# ESCRT-mediated lysosome repair precedes lysophagy and promotes cell survival

**DOI:** 10.1101/313866

**Authors:** Maja Radulovic, Antonino Bongiovanni, Kay O. Schink, Viola Nähse, Eva M. Wenzel, Frank Lafont, Harald Stenmark

## Abstract

Although lysosomes perform a number of essential cellular functions, damaged lysosomes represent a potential hazard to the cell. Such lysosomes are therefore engulfed by autophagic membranes in the process known as lysophagy, which is initiated by recognition of luminal glycoprotein domains by cytosolic lectins such as Galectin-3. Here we show that, under various conditions that cause injury to the lysosome membrane, components of the endosomal sorting complex required for transport (ESCRT) machinery are recruited. This recruitment occurs before that of Galectin-3 and the lysophagy machinery. Subunits of the ESCRT-III complex show a particularly prominent recruitment, which depends on the ESCRT-I component TSG101 and the TSG101- and ESCRT-III-binding protein ALIX. Interference with ESCRT recruitment abolishes lysosome repair and causes otherwise reversible lysosome damage to become cell lethal. Vacuoles containing the intracellular pathogen *Coxiella burnetii* show reversible ESCRT recruitment, and interference with this recruitment reduces intravacuolar bacterial replication. We conclude that the cell is equipped with an endogenous mechanism for lysosome repair which protects against lysosomal damage-induced cell death but which also provides a potential advantage for intracellular pathogens.

## INTRODUCTION

Lysosomes are essential organelles that carry out numerous cellular functions, including degradation of macromolecules, pathogen killing, and signaling functions. On the other hand, because of their low intraluminal pH and high content of Ca^2+^ and enzymes that can potentially trigger cell death, lysosome damage caused by pathogens, sharp crystals, amphiphilic drugs or other membrane-disrupting agents impose a serious threat to cell viability^1, 2^. Previous work has uncovered a lysosome-protective function of heat-shock protein 70^3^, and the existence of an autophagic pathway that sequesters and degrades damaged lysosomes^4, 5^. This pathway, termed lysophagy, is triggered by sensing of lysosomal membrane lesions by cytosolic lectins such as Galectin-3, which recognize exposed intraluminal carbohydrate chains of lysosomal glycoproteins^2, 5^. This is followed by ubiquitination of lysosomal membrane proteins, processing by the ATPase p97, and mobilization of LC3-containing autophagic membranes^2^.

The existence of pathways that protect lysosomes from damage and incapacitate damaged lysosomes has begged the question whether repair mechanisms for damaged lysosomes also exist^2^. Here we have tested the hypothesis that the endosomal sorting complex required for transport (ESCRT) machinery^6^ might play a role in lysosome repair and thereby have a cytoprotective function. Originally identified for its function in protein sorting to the yeast lysosome equivalent, the vacuole^7^, the ESCRT machinery has recently been shown to have a more general function in membrane involution and scission processes that occur in the direction “away” from cytosol, such as virus budding from the plasma membrane, daughter cell separation during cytokinesis, and sealing of the nuclear envelope during mitotic exit^8,9^ Interestingly, the ESCRT machinery has also been shown to mediate repair of both the plasma membrane and the nuclear envelope^10-13^, raising the possibility that it might also function in repair of other membranes.

Based on biochemical and genetic evidence, the ESCRT machinery can be divided into 4 subcomplexes termed ESCRT-0, -I, -II and -III, of which ESCRT-III is thought to mediate membrane sealing/scission through formation of membrane-active oligomeric filaments^8^. Even though ESCRT-0 and -II are important during endosomal protein sorting into intraluminal vesicles of endosomes^14-18^, these subcomplexes appear to be dispensable for most known ESCRT functions^8^. On the other hand, ESCRT-I is frequently involved in recruitment of ESCRT-III, and so is ALIX, a Bro1 domain-containing protein which can potentially form a physical link between ESCRT-I and ESCRT-III^8, 19^.

Here we show that ESCRT-III is indeed recruited to damagedlysosomes, and that this requires ESCRT-I and ALIX. Interference with this mechanism abolishes the cell’s ability to repair damaged lysosomes and causes otherwise reversible lysosome damage to become cell lethal. Surprisingly, we also find that ESCRTs are not only recruited to vacuoles containing the replicating form of the intracellular bacterium *Coxiella burnetii*, but they also provide the bacterium with an advantage by maintaining a niche for its intracellular replication.

## RESULTS

### The ESCRT-III subunit CHMP4B is recruited to damaged lysosomes

In order to achieve permeabilisation of lysosomes in an acute manner we used L-leucyl-L-leucine methyl ester (LLOMe), which is converted into a membranolytic polymeric form in the lysosome lumen by lysosomal hydrolases^20^. Using an antibody against Galectin-3 as marker for lysosome permeabilisation^21^, we observed by confocal fluorescence microscopy that Galectin-3 translocated to vesicular structures in HeLa cells treated with LLOMe, and its colocalisation with the lysosome marker LAMP1 confirmed that these structures are lysosomes (Fig. 1A). We generated a stable HeLa line expressing low levels of mCherry-tagged Galectin-3 and the major ESCRT-III subunit CHMP4B tagged with eGFP and monitored the effect of LLOMe on CHMP4B distribution. Interestingly, whereas both mCherry-Galectin-3 and CHMP4B-eGFP displayed mainly cytosolic staining in untreated cells, treatment with low (250 μM) concentration of LLOMe for 1 hour caused a strong redistribution of both molecules to vesicles positive for the late endosome/lysosome marker CD63 (Fig. 1B). Likewise, antibodies against CHMP4B stained CD63-, Galectin-3- and LAMP1-positive lysosomes after LLOMe incubation of HeLa, RPE1 and H-460 cells, in contrast to untreated cells (Suppl.Fig. 1A-C). We conclude that LLOMe-induced membrane damage causes recruitment of CHMP4B to lysosomes in various cell types.

**Figure 1.**
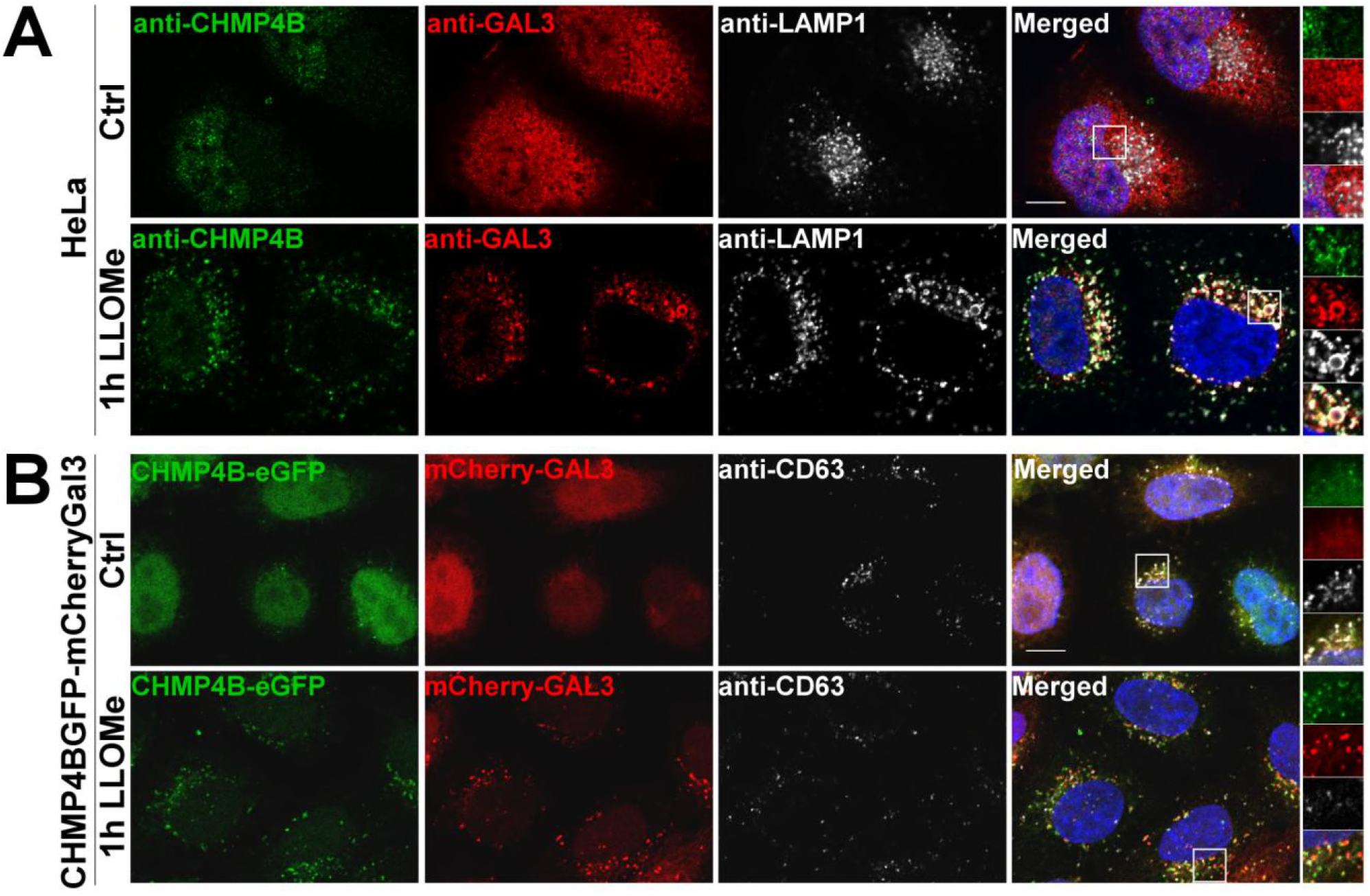
The ESCRT-III subunit CHMP4B is recruited to LAMP1 or CD63 and Gal3-positive damaged endolysosomal membranes. A) Representative confocal images showing recruitment of ESCRT-III complex at the damaged endolysosmal membranes compared to the control. HeLa cells were treated with 250 μM lysosomotropic drug LLOMe or equal volume of DMSO (Ctrl) for 1h before fixation. Endogenous CHMP4B, GAL3 and LAMP1 were immunostained. B) Representative confocal images of HeLa cells stably expressing CHMP4B-eGFP and mCherry-Galectin3 treated and fixed as in A) and stained with CD63 antibody. Scale bars: 10 μm.

We next investigated whether other agents that cause lysosomal membrane damage also induce CHMP4B recruitment. Glycyl-L-phenylalanine-beta-naphthylamide (GPN) is known to cause lysosomal membrane permeabilisation via osmotic swelling^22^, and we indeed observed that this compound caused a profound recruitment of both mCherry-Galectin-3 and CHMP4B-eGFP to LAMP1-containing lysosomes (Suppl. Fig. 2A). Amphiphilic antihistamines have been shown to induce permeabilisation of the lysosome membrane^23^, and both the antihistamines Terfenadine and Astemizole induced recruitment of Galectin-3 and CHMP4B to LAMP1-positive lysosomes, as detected with both fluorescently tagged proteins and antibodies against the endogenous proteins (Suppl.Fig. 2B,C). Collectively, these results show that CHMP4B, like Galectin-3, is recruited to lysosomes upon various types of membrane damage.

### ESCRT-I, ESCRT-III, ALIX and VPS4A are recruited to damaged lysosomes

The observed recruitment of CHMP4B to damaged lysosomes begged the question of which other ESCRT subunits are recruited. Specifically, because ESCRT-0, -I and –II are upstream of ESCRT-III in endosomal sorting^17, 24^, we wondered if this might be the case with recruitment to damaged lysosomes as well. In HeLa cells treated with LLOMe for 2 hours, we found no evidence for recruitment of the ESCRT-0 component HRS or the ESCRT-II subunit VPS36/EAP45. In contrast, the ESCRT-I subunit TSG101 was clearly recruited, as was VPS4A^9^, an ATPase that controls ESCRT-III dynamics (Fig. 2 and Suppl.Fig. 3). As expected, not only CHMP4B, but also two other ESCRT-III subunits, CHMP2A and CHMP3, were found to be recruited, as was the ESCRT-III-related protein IST1 (Fig. 2 and Suppl.Fig. 3). Interestingly, ALIX, a Bro1 domain-containing protein that can bridge ESCRT-I with ESCRT-III^19^, was also recruited, whereas we were unable to detect recruitment of another Bro1 domain protein, HD-PTP (Fig. 2). We were also unable to detect recruitment of the VPS4 isoform VPS4B (Suppl.Fig. 3). From these studies we conclude that ESCRT-I, ESCRT-III, ALIX and VPS4A are recruited to damaged lysosomes.

**Figure 2.**
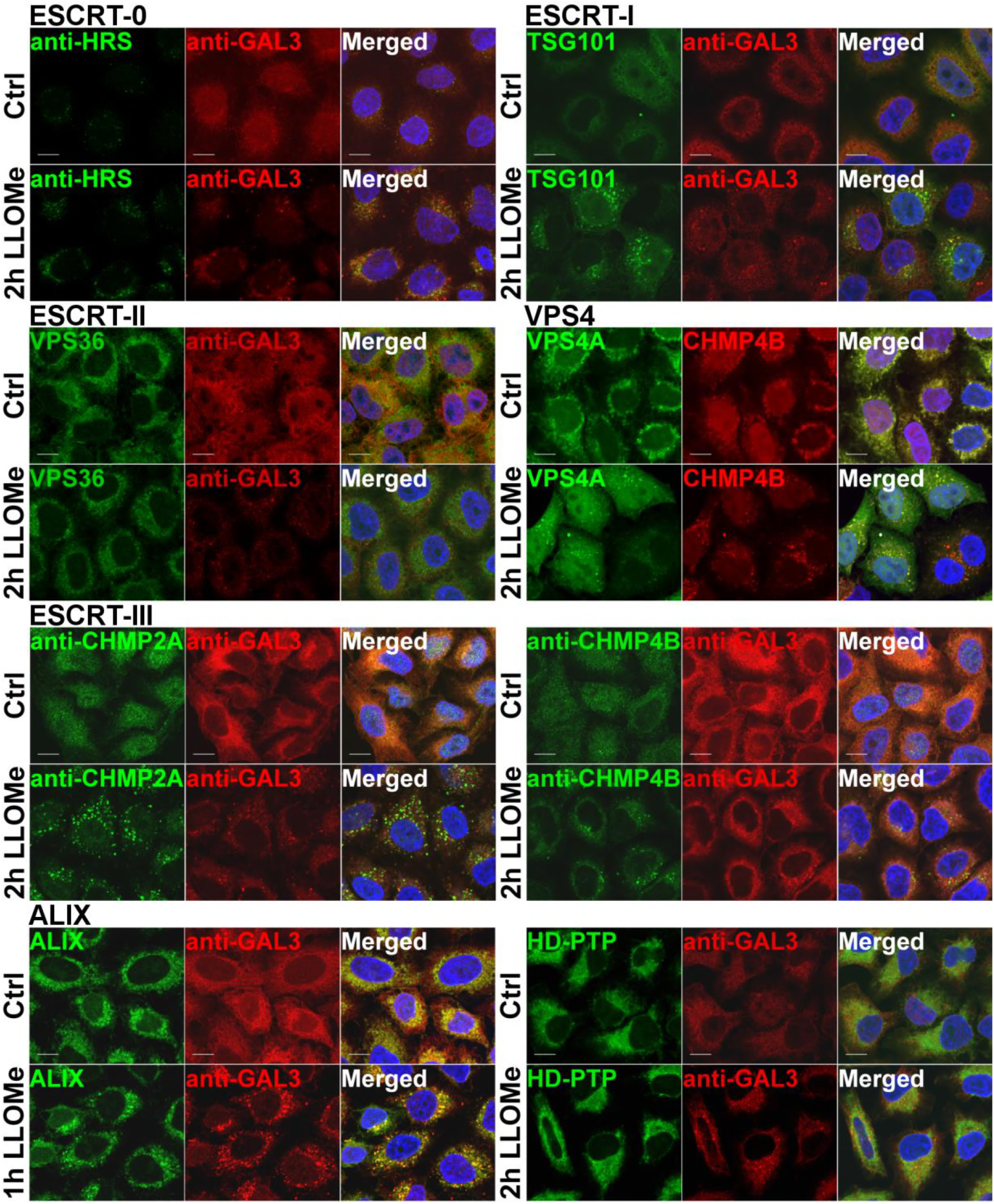
ESCRT-I, ESCRT-III, ALIX and VPS4A are recruited to damaged endolysosomes. In order to screen for ESCRT proteins that are involved in the endolysosomal repair process, cells were incubated with 500 μM LLOMe for 2h and processed for immunofluorescence. ESCRT-0 complex is not involved as HRS does not get recruited to the damaged membranes. TSG101, a component of ESCRT-I complex, is clearly recruited to the sites of damage compared to the DMSO control (Ctrl). The ESCRT-III complex together with ALIX is recruited to the damaged endolysosomes. In contrast, ESCRT-II and HD-PTP show no recruitment. Scale bars: 10 μm.

### TSG101 depletion inhibits CHMP4B recruitment on damaged lysosomes whereas CHMP2A knockdown stabilizes it

The above results suggested that TSG101 and ALIX might have a role in ESCRT-III recruitment, and we tested this further using siRNA to deplete various ESCRT proteins in HeLa cells expressing CHMP4B-eGFP and monitoring CHMP4B-eGFP recruitment by live microscopy. Knock-down of the ESCRT-0 subunit HRS was without effect on CHMP4B-eGFP recruitment to lysosomes upon LLOMe treatment of the cells (Fig. 3A, Suppl. Movie S1), consistent with our finding that this ESCRT subunit is not recruited itself. On the other hand, depletion of TSG101 caused a strong delay in CHMP4B-eGFP recruitment (Fig. 3B, Suppl.Movie S2). Conversely, depletion of CHMP2A caused increased accumulation of CHMP4B-eGFP on LLOMe-damaged lysosomes (Fig. 3C, Suppl. Movie S3), in agreement with previous studies on ESCRT recruitment to other membranes, suggesting that CHMP2A limits the extent of CHMP4B recruitment^25^. Whereas depletion of ALIX was without detectable effect on CHM4B-eGFP recruitment (Fig. 3C, Suppl.Movie S3), co-depletion of TSG101 and ALIX led to an almost complete lack of CHMP4B-GFP recruitment (Fig. 3D, Suppl.Movie S4). These results confirm that TSG101 in ESCRT-I is upstream of ESCRT-III in recruitment to damaged lysosomes and also reveal a role for ALIX.

**Figure 3.**
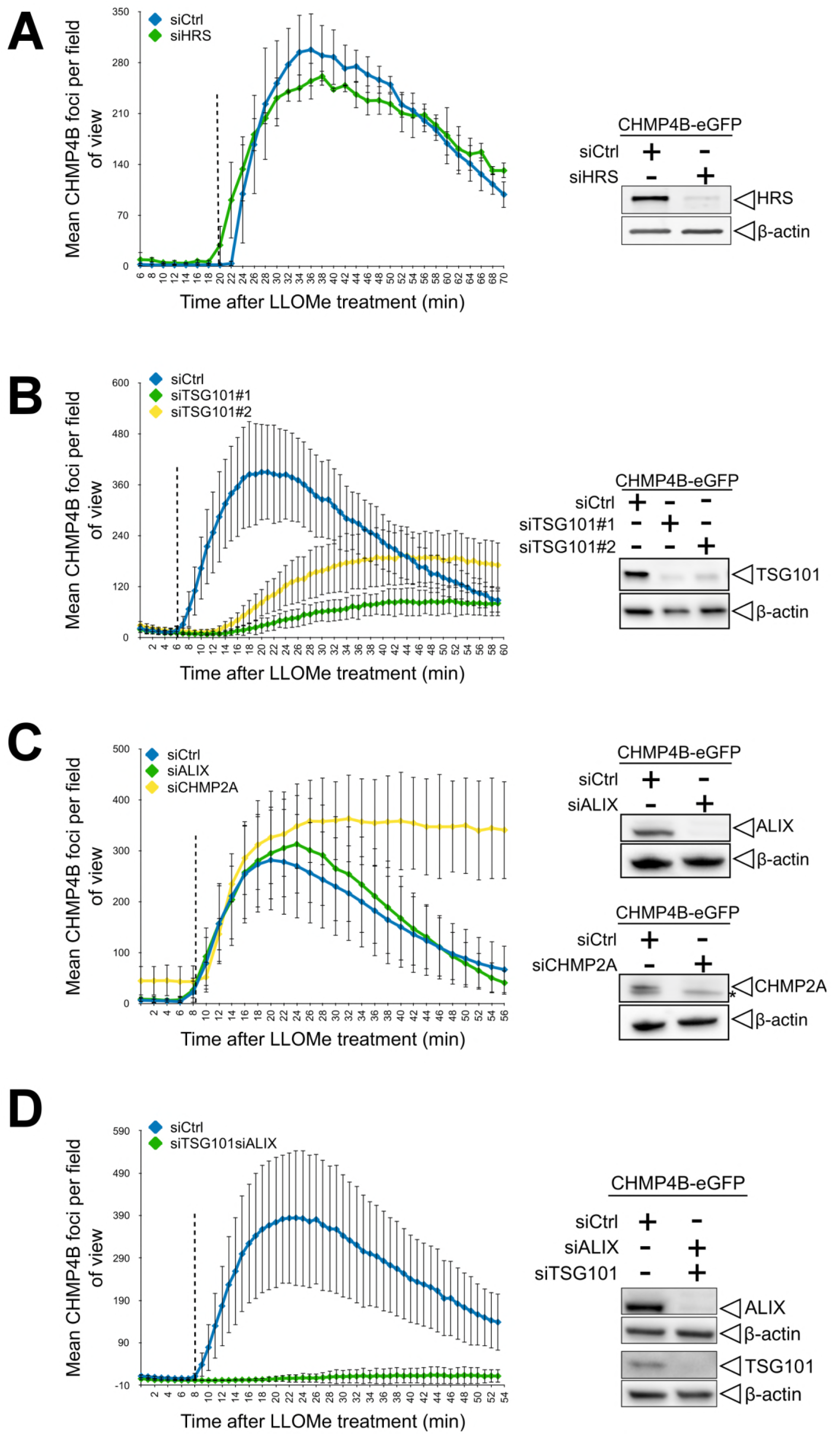
Dynamics of CHMP4B accumulation at the damaged endolysosomal membranes. HeLa cells stably expressing CHMP4B-eGFP were transfected with siRNAs against: (A) HRS, (B) TSG101, (C) ALIX, and (D) both TSG101 and ALIX. A) Depletion of HRS did not alter the dynamics of CHMP4B-eGFP as compared to siCtrl during treatment with LLOMe. B) Downregulation of TSG101, with two independent siRNAs cause a delay in CHMP4B-eGFP recruitment as compared to siCtrl, indicating an important role of ESCRT-I in recruiting the downstream components. C) siRNA mediated depletion of ALIX shows no significant change, while CHMP2A depletion increases CHMP4B-eGFP positive foci. D) Simultaneous depletion of TSG101 and ALIX caused almost no recruitment of CHMP4B upon induction of endolysosomal damage. Left panel shows quantification graphs for each condition (between 10 and 15 fields of view representing more than 200 cells per condition were analyzed; error bars indicate SD) where dotted line indicates when LLOMe was added to the cells. Right panel shows knockdown efficiency of siRNA oligos as detected by Western blot (*, nonspecific immunoreactivity).

### ESCRT is required for repair of the lysosome membrane

We next asked whether the ESCRT machinery is involved in repair of damaged lysosome membranes. As an assay for membrane permeability we used the ability of lysosomes to retain LysoTracker^®^, a weak base which accumulates in acidic lysosomes and is fluorescent at low pH^26^. As expected, incubation of HeLa cells with LLOMe at 250 μM caused reduced LysoTracker staining of lysosomes (Fig. 4A). Interestingly, however, at this relatively low LLOMe concentration, LysoTracker staining fully recovered within 30-60 minutes, indicating that the LLOMe-induced membrane damage was repaired (Fig. 4A). Importantly, whereas LysoTracker staining recovered after LLOMe treatment of control cells, lysosomes in cells depleted for TSG101 and ALIX failed to recover LysoTracker fluorescence (Fig 4C). This indicates that the ESCRT machinery mediates repair of damaged lysosomes.

**Figure 4.**
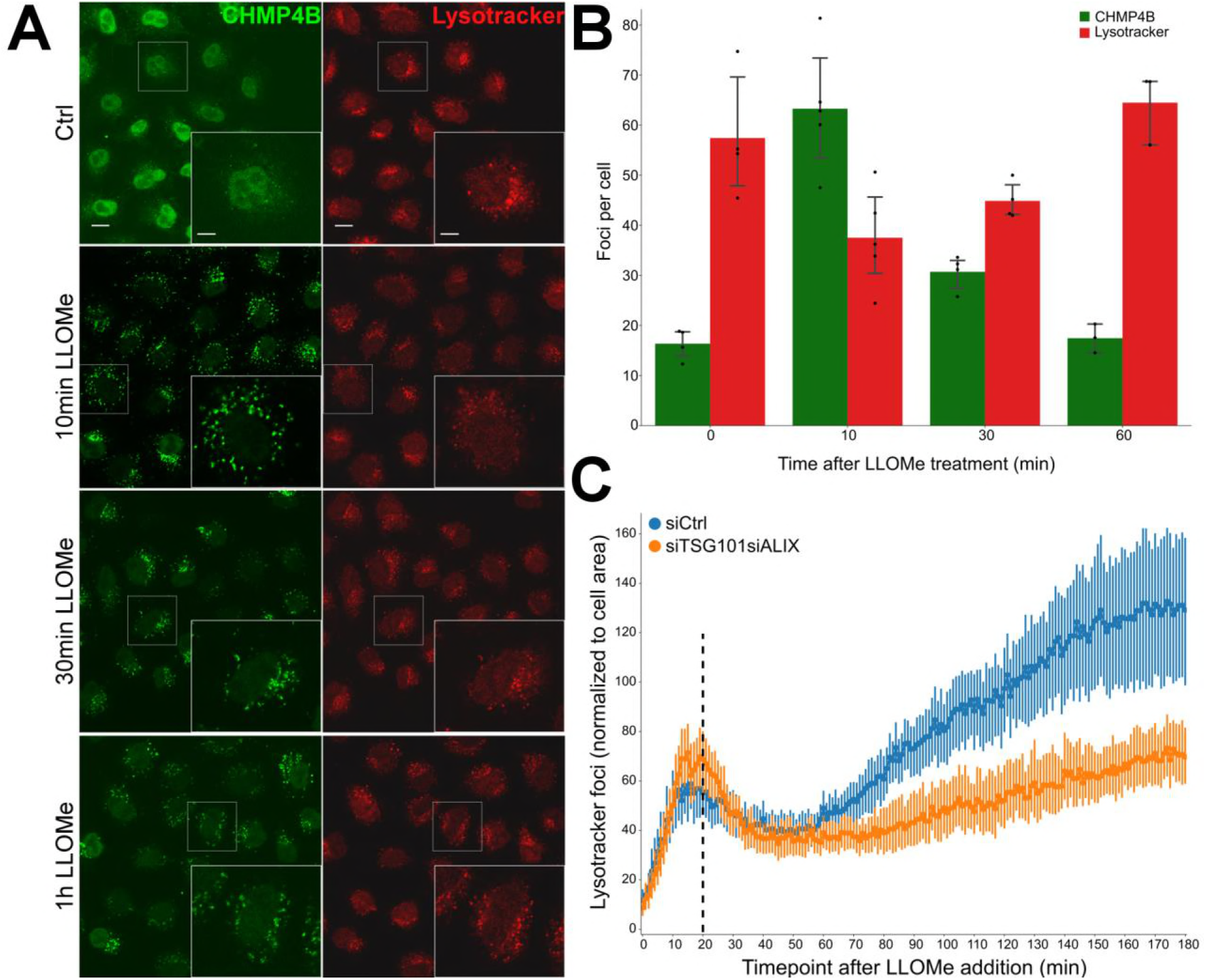
ESCRTs are essential for the repair after endolysosomal damage. A) HeLa cells stably expressing CHMP4B-eGFP were treated with 250 μM LLOMe and 75 nM Lysotracker DND-99 and fixed at different time points as indicated. After 10 min of LLOMe treatment CHMP4B is recruited whereas the number of lysotracker positive foci is reduced. After 30 min, lysosomes gain back functionality (judging by the increased number of Lysotracker foci) and appear fully recovered after 1h indicating that the ESCRT complex is able to seal the damaged endolysosomal membranes. B) Quantification graph showing CHMP4B and Lysotracker positive foci per cell at different time points. C) HeLa cells stably expressing CHMP4B-eGFP were co-transfected with siRNAs against TSG101 and ALIX or siCtrl. 48h post-transfection, cells were pre-treated for 20 min with 75nM Lysotracker Deep-Red, which was used as a read-out. Dotted line indicates when LLOMe was added to the cells pre-incubated with Lysotraker. While the decrease in the number of lysotracker spots is quickly recovered in the control (siCtrl), simultaneous depletion of ALIX and TSG101 led to a severe impairment in lysosomal repair. Graph is normalized to the area occupied by the cells.

### ESCRT-III shows much faster recruitment kinetics than Galectin-3 to damaged lysosomes

In order to monitor the dynamics of ESCRT recruitment to damaged lysosomes, we used live microscopy to study HeLa cells stably expressing low levels of CHMP4B-eGFP and mCherry-Galectin-3. Upon LLOMe addition, mCherry-Galectin-3 accumulated slowly on the lysosomes as expected (Fig. 5). Remarkably, however, CHMP4B-eGFP accumulated much faster, with detectable levels already after 1 minute and reaching a peak level at about 10 minutes (Fig. 5 and Suppl. Movie S5). This indicates that the ESCRT machinery detects a subtler membrane damage than the exposure of intraluminal β-galactosides detected by Galectin-3.

**Figure 5.**
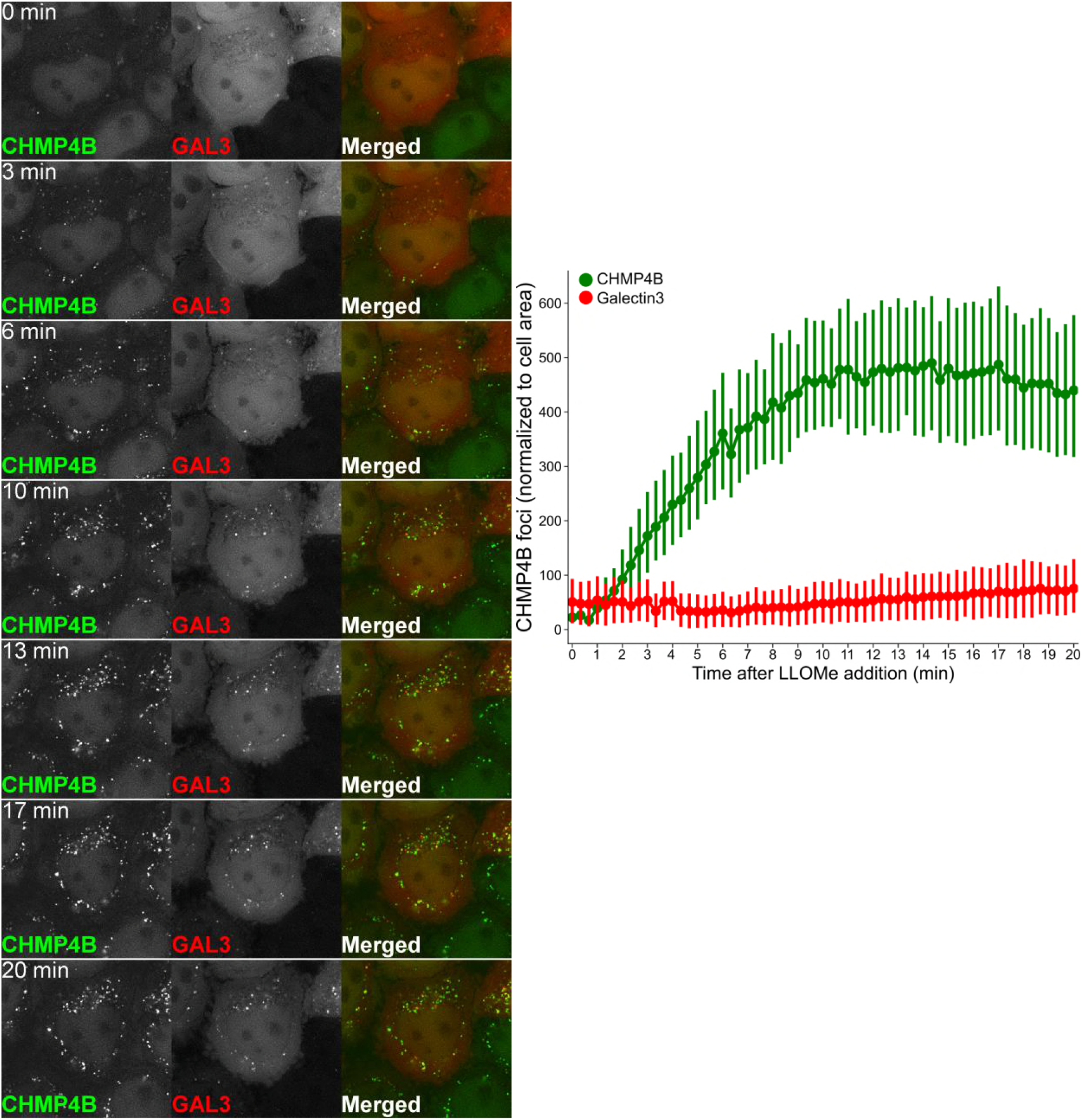
Dynamics of CHMP4B recruitment to the damaged endolysosomal membranes. Representative movie montage from live cell imaging comparing the timing of CHMP4B-eGFP and mCherry-Galectin3 recruitment. Using the perfusion system, 250 μM LLOMe was added to HeLa cells stably expressing CHMP4B-eGFP and mCherry-Galectin3 at 20 sec while imaging. As shown in the representative movie montage and the quantification graph, CHMP4B recruitment precedes Galectin3, which is an established marker for detection of damaged endolysosomal membranes. Quantification graph (right panel) is normalized to the area occupied by the cells, at least 40 cells.

### ESCRT-III recruitment to damaged lysosomes precedes lysophagy

Because Galectin-3 recruitment to lysosomes triggers lysophagy, we next compared the dynamics of CHMP4B recruitment with the dynamics of various molecules involved in lysophagy by confocal microscopy of HeLa cells fixed at various time points after LLOMe addition. Ubiquitination of lysosomal membrane proteins is known to occur early during lysophagy^5^, and we therefore monitored the kinetics of ubiquitin acquisition on lysosomes using an antibody against conjugated ubiquitin. Like with Galectin-3 recruitment, ubiquitin appeared on the damaged lysosomes much slower than CHMP4B (Fig. 6A and Suppl.Fig. 4A). We also studied recruitment of the autophagy adaptor p62/SQSTM1, a protein known to connect ubiquitinated autophagic cargoes to autophagic membranes^5, 27^, and a major constituent of the autophagy machinery, LC3^5, 28^. Both p62 and LC3 appeared on damaged lysosomes with much slower kinetics than CHMP4B (Fig. 6B and Suppl.Fig. 4B). The autophagy inhibitor SAR405^29^ was without effect on CHMP4B-GFP recruitment (Suppl.Movie S6). We conclude from these studies that the ESCRT machinery is recruited much prior to the autophagy machinery to damaged endosomes.

**Figure 6.**
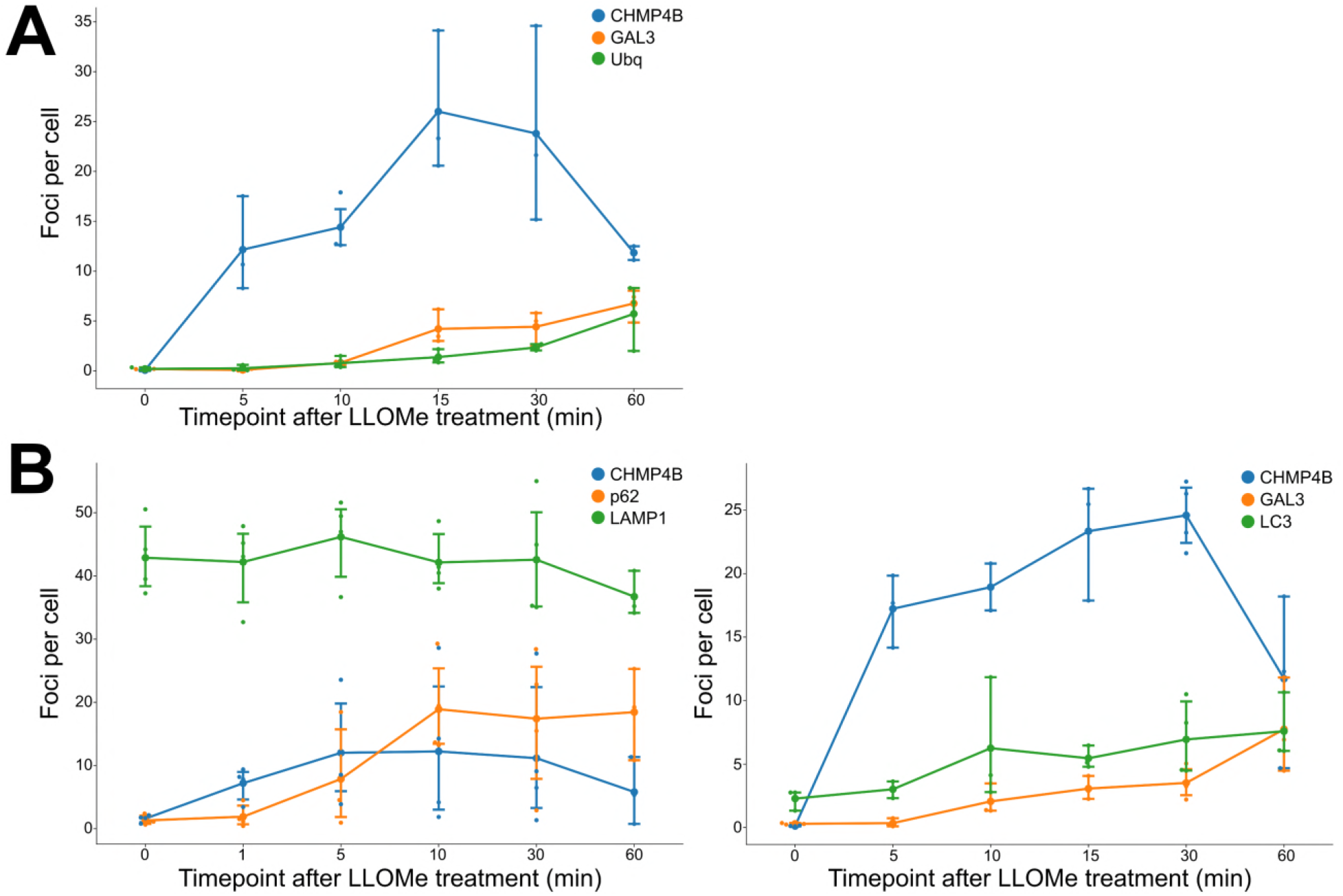
ESCRT-III recruitment to damaged lysosomes precedes lysophagy. HeLa cells stably expressing CHMP4B-eGFP and/or mCherry-Galectin3 were incubated with 250 μM LLOMe, fixed at different time points as indicated and stained for different markers. A) CHMP4B is recruited before ubiquitin and Galectin-3 upon lysosomal membrane damage. B) CHMP4B is recruited before p62 and LC3 on the damaged lysosomes. Dynamics of LAMP1-positive foci appears stable independently of LLOMe treatment. Quantification graphs are normalized to the number of foci per cell, at least 100 cells per condition.

### ESCRT recruitment protects cells against cell death caused by lysosome damage

Having found that ESCRT recruitment mediates lysosome repair and occurs prior to lysophagy, we next addressed the biological importance of this mechanism. By live imaging of TSG101-ALIX-depleted cells treated with LLOMe, we had observed that some cells started dying upon prolonged incubation (Suppl.Movie S4), and we therefore hypothesized that ESCRT-mediated lysosome repair might provide a cell survival advantage. To test this hypothesis, we depleted cells of TSG101, ALIX, or both, and used flow cytometry to monitor cell death 6 hours after LLOMe addition. Strikingly, whereas LLOMe at 250 μM had minimal effect on cell viability, siRNA-mediated depletion of TSG101 led to a dramatic rise in cell death in the presence of LLOMe. While depletion of ALIX had only a minor effect on cell death, the combined depletion of both TSG101 and ALIX led to death of a large fraction of the cells (Fig. 7, Suppl.Fig. 5, and Suppl.Movie S7). This indicates that ESCRT-mediated lysosome repair plays an important role in promoting cell viability after lysosome injury.

**Figure 7.**
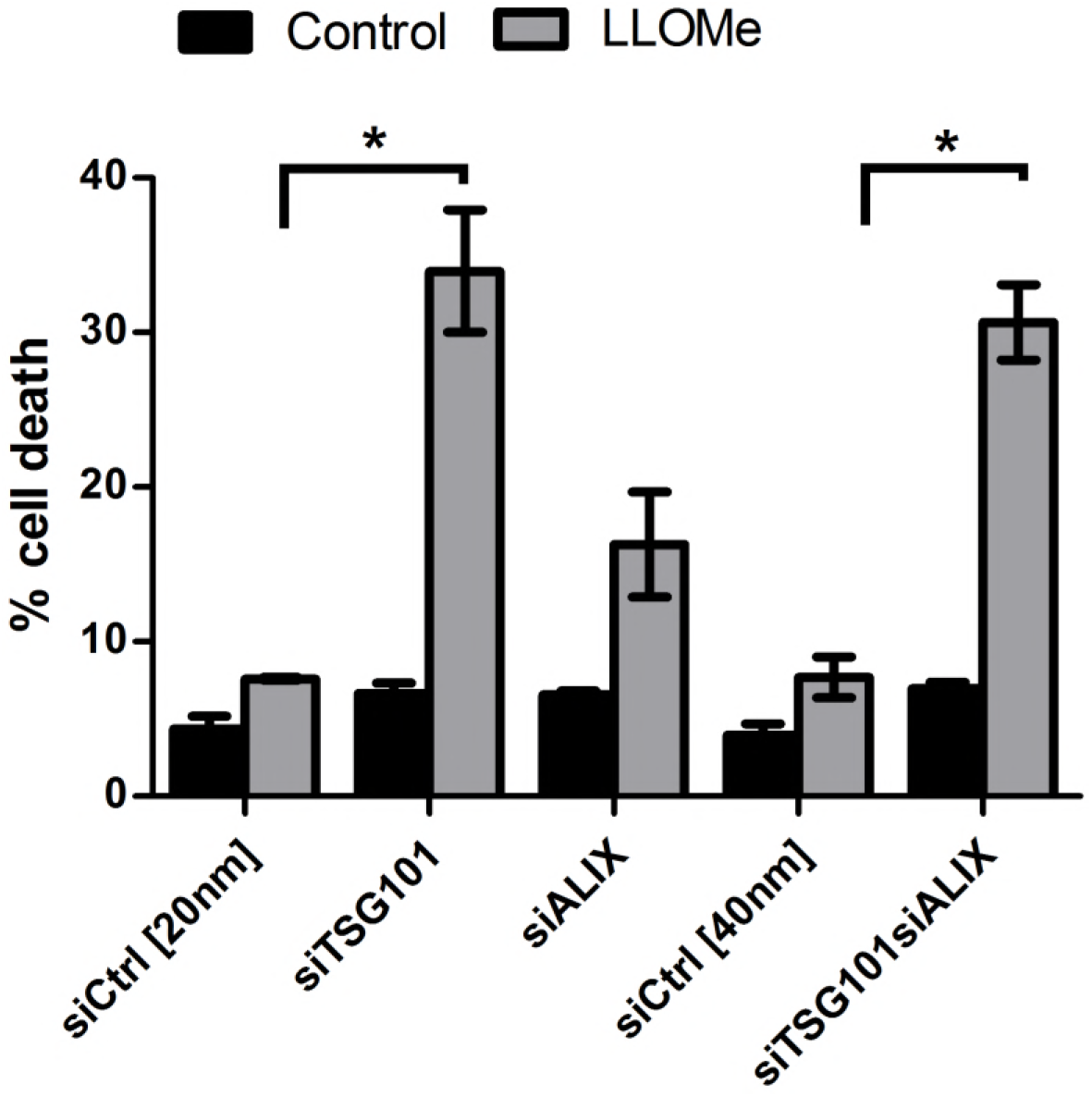
ESCRTs are essential for cell viability after endolysosomal damage. Cell viability in cells after siRNA mediated depletion of TSG101 and ALIX. HeLa cells stably expressing CHMP4B-eGFP were transfected with different siRNAs as indicated. Treatment of TSG101 or TSG101+ALIX siRNA transfected cells with 250 μM LLOMe for 6h showed increased cell death compared to the control as measured by flow cytometry. Error bars show SEM of three independent experiments. Statistical significance was determined using Student’s t-test where * indicates p-value ≤ 0.05. p-values are presented only when significant compared to siCtrl.

### ESCRT recruitment to *Coxiella burnetii* vacuoles promotes bacterial replication

Many intracellular pathogens are able to survive and even replicate within modified phagosomes of host cells^30^. One very interesting example is the small Gram-negative bacterium *Coxiella burnetii*, the causative agent of Q fever^31^. *C.burnetii* is an obligate intracellular pathogen which has the remarkable property of replicating inside an acidic lysosome-like vacuole. A recent study has revealed that Galectin-3 is recruited to *C.burnetii* vacuoles^32^, and we therefore asked whether the ESCRT machinery might be recruited as well. To address this question, we infected HeLa cells expressing mCherry-Galectin-3 and CHMP4B-eGFP with *C.burnetii* and used long-time live microscopy to monitor protein recruitment. Interestingly, after a lag time of several hours, bacterium-containing vacuoles became positive for both Galectin-3 and CHMP4B before they quickly turned negative again. This was repeated several times (Fig. 8A and Suppl. Movie S8), and the vacuole expansion that occurred after CHMP4B and Galectin-3 recruitment indicated that the replicative niche was kept intact (Suppl.Movie S9). We interpret this as sporadic ruptures of the vacuole membrane which were repeatedly repaired by the ESCRT machinery.

**Figure 8.**
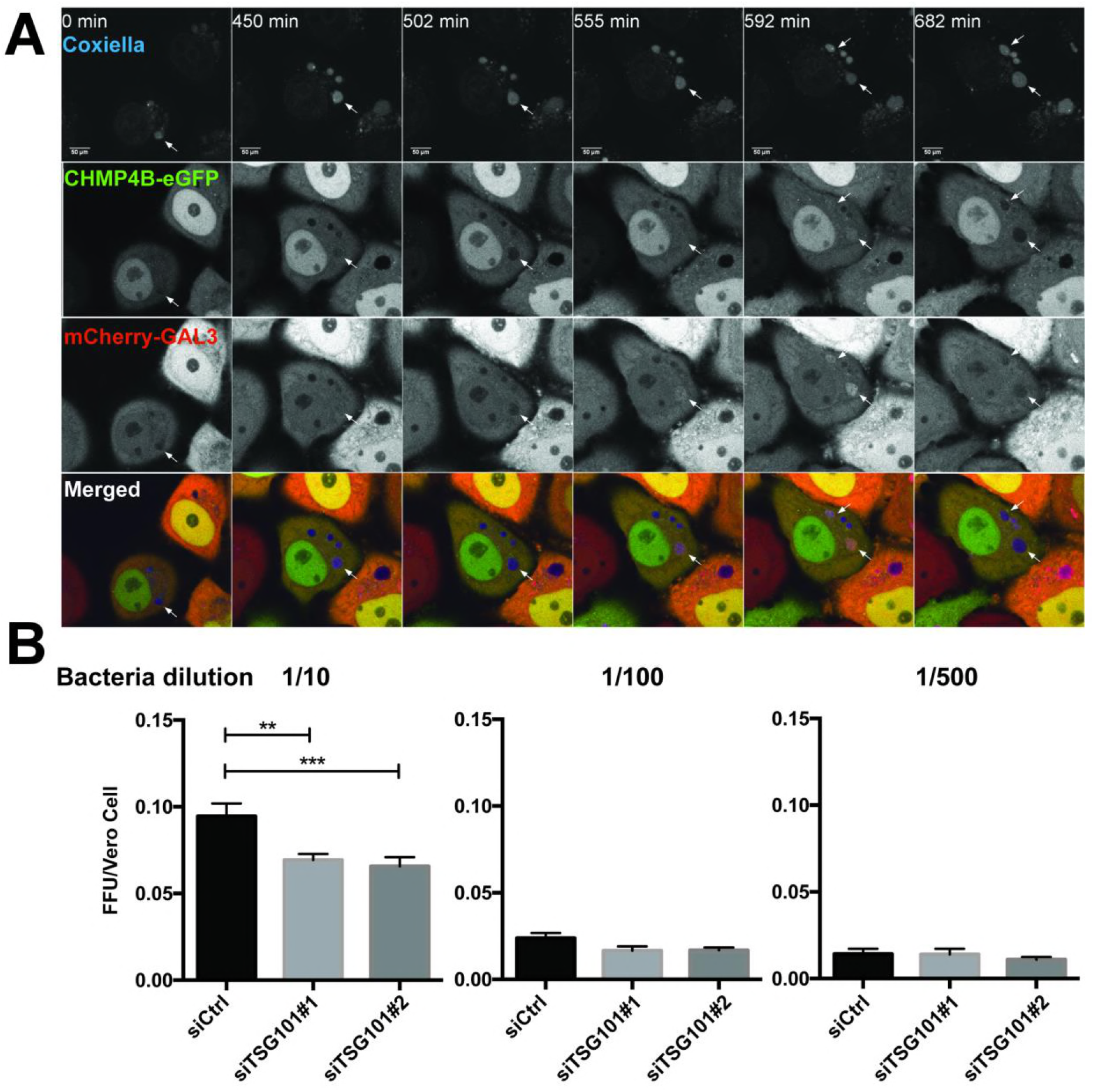
ESCRT-III and Galectin-3 are recruited to the *Coxiella burnetii* vacuole. A) *Coxiella*-containing vacuole becomes positive for CHMP4B and Galectin-3. HeLa cells stably expressing CHMP4B-eGFP were transfected with the mCherry-Galectin-3 plasmid. 24h later, cells were infected with WT *Coxiella burnetti*. Live cell imaging was performed on a spinning disk module. Recording started 24h post-infection and several time points are indicated during the next 24h of observation (time indicated in minutes, see Suppl.Movie S8). Arrows indicate *Coxiella*-containing vacuole (DRAQ5 labeling) becoming positive for CHMP4B-eGFP and mCherry-Galectin-3. B) Bacterial viability and replication assay upon TSG101 KD. HeLa cells were treated with either control siRNA or two different TSG101 siRNAs for 48h. Then cells were infected with mCherry-*Coxiella burnetii* for 48h before lysis and serial dilutions were made to infect Vero cells. 72h later, infected cells were fixed, DAPI stained and processed for quantitative image analysis. Between 30 and 40 fields representing more than 9,000 cells per condition of 3 independent experiments were analyzed. Statistical significance was determined using one-way ANOVA test. **p-value≤ 0.01, ***p-value≤ 0.001.

We hypothesized that failure of ESCRT-mediated vacuole repair might interfere with bacterial infection. To test this hypothesis, we performed a Fluorescent Foci Unit (FFU) assay in which we used lysates from *C.burnetii*-containing vacuoles from HeLa cells with or without depletion of TSG101 to infect Vero cells at different dilutions. Interestingly, the bacterial viability and replication FFU assay showed that depletion of TSG101 decreased *C.burnetii* replication ( Fig. 8B). We conclude that the ESCRT machinery provides the bacterium a replicative advantage which can probably be explained by the requirement of an intact vacuole for efficient *C.burnetii* replication to proceed.

## DISCUSSION

Here we have shown that ESCRT-mediated repair of damaged lysosomes occurs independently of lysophagy, which is in excellent agreement with results reported in a very recent paper^33^. In addition, we show that this mechanism is crucial for promoting cell viability after lysosome injury and that ESCRT-mediated membrane repair can also be exploited for maintaining an intact replicative niche for an intracellular pathogen. Thus, ESCRT-mediated repair of lysosomal membranes can have both beneficial and harmful consequences for the host cell.

Whereas lysophagy is triggered by holes in the lysosome membrane large enough for detection of intraluminal β-galactosides by cytosolic lectins such as Galectin-3, the ESCRT machinery must be recruited by subtler cues. Because knockdown of the ESCRT-I subunit TSG101 resulted in a strong delay in ESCRT-III recruitment, ESCRT-I is likely to mediate recruitment of ESCRT-III to sites of injury in the lysosomes membrane. Even though depletion of ALIX by itself had no detectable effect on ESCRT-III recruitment, simultaneous depletion of ALIX and TSG101 resulted in a complete block of ESCRT-III recruitment, suggesting that ALIX and TSG101 cooperate to recruit ESCRT-III. This resembles the two-pronged recruitment of ESCRT-III by ESCRT-I and Bro1/ALIX during endosomal sorting and cytokinetic abscission^34, 35^.

Previous work has uncovered cellular mechanisms that protect the lysosome membrane from damage^3^, and that lead to elimination of damaged lysosomes^4, 5^. We have here shown the existence of an ESCRT-mediated lysosomal repair pathway that kicks in before engagement of the lysophagy machinery. Thus, the cell is equipped with several strategies for preventing leakage of lysosomal contents into cytosol. Our finding that otherwise reversible lysosomal membrane damage becomes cell lethal in the absence of ESCRT recruitment illustrates the importance of the lysosomal repair pathway.

Many intracellular pathogens are able to replicate inside host cells, and in some cases replication can take place in lysosome-like bacterial vacuoles. *C.burnetii* is a good example of this^31^, and it was interesting to observe the reversible recruitment of ESCRT proteins to the *C.burnetii* vacuole. Galectin-3 was also recruited, although our time-lapse movies had too low temporal resolution to determine whether Galectin-3 was recruited after ESCRT-III, as found with damaged lysosomes. The fact that both ESCRT-III and Galectin-3 recruitment was reversible suggests that the *C.burnetii* vacuoles undergo sporadic membrane damage that can be repaired by the ESCRT machinery. Upon recruitment and repair, Galectin-3 might be degraded in the lysosome-like lumen of the vacuole whereas ESCRT-III would dissociate upon fulfilling its function in membrane repair. Our finding that ESCRT depletion inhibits *C.burnetii* replication indicates that ESCRT-mediated repair of the sporadically injured vacuole membrane is required for keeping the vacuole intact over the several hours required for optimal replication.

Our findings raise several questions and perspectives. First, how is lysosomal membrane repair achieved by ESCRT proteins? This has to be clarified by future experiments, but our working hypothesis is that the membrane patch containing the lesion becomes internalized into intraluminal vesicles by the same mechanism as that used in ESCRT-mediated endosomal protein sorting and biogenesis of multivesicular endosomes^36^. Another important question concerns the cues that trigger ESCRT recruitment to damaged lysosomes. Ca^2+^ leakage out of the injured lysosome, detected by ALIX and its Ca^2+^ binding partner ALG2, has been put forward as a mechanism^33^, but we note that TSG101 appears to be more important than ALIX for mediating ESCRT-III recruitment. We therefore hypothesize that additional mechanisms may exist for ESCRT recruitment to damaged lysosomes. Lysosome damage has been shown to induce cell death by several alternative mechanisms^2^, and it will be interesting to learn which principal mechanism causes cell death upon lysosome damage in ESCRT-depleted cells. Our observation that the ESCRT machinery facilitates intracellular replication of *C.burnetii* raises the question whether other intracellular pathogens also take advantage of the ESCRTs. If so, pharmacological ESCRT inhibition might offer a potential strategy for future treatment of infections caused by intracellular pathogens.

## MATERIALS AND METHODS

### Reagents, cell culture and stable cell lines

Lysosomotropic drugs used in this study, L-Leucyl-L-Leucine methyl ester (cat. no. 16008) and Gly-Phe-β-naphthylamide (cat. no. 14634), were purchased from Cayman Chemical and used at 250 μM unless otherwise indicated. The antihistamines Astemizole (cat. no. 2861) and Terfenadine (cat. no. 9652), were purchased from Sigma Aldrich. All compounds were dissolved in dimethyl sulfoxide (DMSO) and stored at -80°C. Other reagents used in this study: Lysotracker DND-99 (cat. no. 7528) for immunoflorescence and Lysotracker DeepRed (cat. no. 12492,) for live cell imaging were purchased from Molecular Probes and were used at 75 nM.

Human HeLa "Kyoto" (HeLaK) and hTERT-RPE-1 cells (human retinal pigment epithelial cells immortalised with telomerase) were maintained in DMEM (Gibco) and F12:DMEM medium, respectively, supplemented with 10% fetal bovine serum (FBS), 5 U/ml penicillin and 50 μg/ml streptomycin. The human large-cell lung carcinoma cell line NCI-H460 was maintained in medium recommended by ATCC. All cells were maintained at 37°C supplemented with 5% CO_2_. A stable HeLa cell line expressing CHMP4B-eGFP was obtained from Anthony A. Hyman (Max Planck Institute for Molecular Cell Biology and Genetics, Dresden, Germany;^37^).

All other stable cell lines used in this study were lentivirus-generated pools, using plasmids pCDH-PGK-IRES-Blast-mCherry-Galectin3 and pCDH-PGK-IRES-Blast-mCherry-ALIX for stable expression of mCherry-Galectin3 and mCherry-ALIX, respectively. The weak PGK promoter was used for transgene expression in order to achieve low expression levels. Third generation lentivirus was generated as previously published in^38^. Briefly, mCherry or eGFP fusions were generated as Gateway ENTRY plasmids using standard molecular biology techniques. From these vectors, lentiviral transfer vectors were generated by recombination into pLenti Destination vectors using a Gateway LR reaction. VSV-G pseudotyped lentiviral particles were packaged using a third-generation packaging system^39^. Cells were then transduced with low virus titers and stable expressing populations were generated by antibiotic selection. Other cell lines stably expressing HRS, TSG101, VPS4A, VPS4B, VPS36, HD-PTP and IST1 were kindly provided by Eva M. Wenzel (Oslo University Hospital) and Coen Campsteijn (University of Oslo, Norway).

### siRNA transfections

Silencer Select siRNAs against HRS (5'-CGACAAGAACCCACACGTC-3'), TSG101 (#1, 5'-CCGUUUAGAUCAAGAAGUA-3'; #2, 5'-CCUCCAGUCUUCUCUCGUC-3'), siCHMP2A (5'-AAGAUGAAGAGGAGAGUGA-3'), ALIX (#1, 5'-GCAGUGAGGUUGUAAAUGU-3'; #2, 5'-CCUGGAUAAUGAUGAAGGA-3'), and nontargeting control siRNA (predesigned, cat. no.4390844) were purchased from Ambion. Cells at 50% confluency were transfected with 20–50 nM final siRNA concentration using Lipofectamine RNAiMax transfection reagent (Life Technologies) according to the manufacturer’s instructions and harvested after 48h.

### Antibodies

Rabbit anti-CHMP3, rabbit anti-ALIX and rabbit anti-CHMP4B antibodies were described previously^35^. Goat anti-Galectin3 antibody (cat. no. AF1154) was purchased from R&D Systems, rabbit anti-CHMP2A (cat. no. 10477-1-AP) from Proteintech. Mouse anti–β-actin (Sigma-Aldrich), mouse anti-GFP (Roche), mouse anti-TSG101 (BD Transduction Laboratories), rabbit anti-IST1 (Proteintech), and goat anti-mCherry (Acris) were used as primary antibodies. Secondary antibodies included anti-mouse, anti-rabbit, and anti-goat Alexa Fluor 488 (Jackson ImmunoResearch), Alexa Fluor 568 (Molecular Probes), Alexa Fluor 647 (Jackson ImmunoResearch).

### Immunoblotting

Cells were lysed in 2X sample buffer (125 mM Tris-HCl, pH 6.8, 4% SDS, 20% glycerol, 200 mM DTT, and 0.004% bromophenol blue). Whole-cell lysates were subjected to SDS-PAGE on 4–20% gradient gels (mini-PROTEAN TGX; Bio-Rad). Proteins were transferred to Immobilon-P membranes (Millipore) followed by blocking and antibody incubation in 5% fat-free milk powder in Tris-buffered saline with 0.01% Tween 20. Membranes incubated with horseradish peroxidase–conjugated antibodies were developed using Clarity Western ECL substrate solutions (Bio-Rad) with a ChemiDoc XRS+ imaging system (Bio-Rad).

### Immunofluorescence staining and confocal fluorescence microscopy

Cells seeded on coverslips were fixed in 4% EM grade paraformaldehyde for 10 minutes, permeabilized with 0.05% saponin in PBS buffer and labeled with antibodies. For HRS localization experiments, cells were permeabilized with PEM buffer (80 mM K-Pipes, pH 6.8, 5 mM EGTA, and 1mM MgCl_2_) before fixation. Stained coverslips were examined with a Zeiss LSM 780 confocal microscope (Carl Zeiss) equipped with an Ar laser multiline (458/488/514 nm), a DPSS-561 10 (561 nm), a laser diode 405-30 CW (405 nm), and a HeNe laser (633 nm). The objective used was a Zeiss Plan-Apochromat 63×/1.40 Oil DIC M27 (Carl Zeiss). Image processing was performed with ImageJ software (National Institutes of Health). Intensity settings for the relevant channels were kept constant during imaging. Images shown in figures are representative of at least three independent experiments.

### Live cell imaging

Cells seeded in MatTek 35mm petri dish, 20 mm Microwell No. 1.5 coverglass were imaged on a Deltavision microscope (Applied Precision) equipped with Elite TruLight Illumination System, a CoolSNAP HQ2 camera and a 60× Plan Apochromat (1.42 NA) lens. For temperature control during live observation, the microscope stage was kept at 37 °C by a temperature-controlled incubation chamber. Time-lapse images (6 *z*-sections 2,2 μm apart) were acquired every 1-3 minutes over a total time period of 4 h, and deconvolved using the softWoRx software (Applied Precision). In addition, Deltavision OMX V4 microscope equipped with three PCO.edge sCMOS cameras, a solid-state light source, a 60x 1.42 NA objective and a laser-based autofocus was used. Environmental control was provided by a heated stage and an objective heater (20-20 Technologies). Images were deconvolved using softWoRx software and processed in ImageJ/FIJI.

### Flow cytometry

Cells were fixed in 70% ethanol and DNA was stained with Hoechst 33342 (1,5 μg/ml). Flow cytometry analysis was performed on LSRII flow cytometer (BD Biosciences) using FACS Diva software. Cell doublets were excluded by gating, and the subG1 fraction was determined by gating cells with a Hoechst signal less than 2N DNA content. Graphs shown in figures are representative of at least three independent experiments.

### *Coxiella* infection

Propagation of mCherry-*Coxiella burnetii* (generously provided by Dr. Robert Heinzen, Rocky Mountain Laboratories, NIAID, NIH, Hamilton, MT, USA) was performed in Vero cells as previously described^40^. A fluorescent foci unit (FFU) infection assay was used to quantify the replication and viability of *C. burnetii* in HeLa cells in 24-well plates treated with control or TSG101 siRNA for 48 h (with silencing extent verified by western blotting). In brief, after 48 h treatment with siRNAs, cells were infected as previously described^41^ for 48 h and lysed. Samples were serially diluted and used to infect Vero cells in a 24 well-plate. After 72 h of infection, Vero cells were fixed, DAPI stained and processed for fluorescent microscopy. Several thousands of cells were automatically scored per condition to determine an average number of FFU for each sample and three independent experiments were pooled.

### Image analysis and post-processing

Images were analyzed in Fiji/ImageJ using custom python scripts. Briefly, images were filtered to remove noise and foci of ESCRTs and/or associated proteins were segmented by the “Find maxima” function and scored. Alternatively, foci were segmented by thresholding and objects were then identified and scored by the “Analyze Particles” function of ImageJ. All postprocessing was performed using Python and the “pandas“ package, plots were generated using Seaborn. For live cell movies, foci counts were normalized to the area occupied by cells within a given field of view. For fixed cell analysis, nuclei were segmented and counted, and all foci counts were normalized to the number of nuclei within a given field of view. Analysis scripts have been deposited at https://github.com/koschink/Radulovic_et_al

## ACKNOWLEDGEMENTS

We thank Coen Campsteijn for kindly providing plasmid constructs, and M.E. Mansilla and M.-I. Colombo for sharing protocols and reagents. M.R. is a postdoctoral fellow of the South-Eastern Norway Regional Health Authority. K.O.S. holds a career development fellowship and V.N. a postdoctoral fellowship from the Norwegian Cancer Society. E.M.W. is a senior research fellow of the South-Eastern Norway Regional Health Authority (grant number 2015014). H.S. is supported by project grants from the Norwegian Cancer Society and the South-Eastern Norway Regional Health Authority. F.L. is supported by the ANR, project number 15-CE15-0017. This work was partly supported by the Research Council of Norway through its Centres of Excellence funding scheme, project number 262652.

